# Gut microbial diversity in stingless bees is linked to host wing size and is influenced by geography

**DOI:** 10.1101/2021.07.04.451070

**Authors:** Hongwei Liu, Mark A. Hall, Laura E. Brettell, Megan Halcroft, Juntao Wang, Scott Nacko, Robert Spooner-Hart, James M Cook, Markus Riegler, Brajesh Singh

## Abstract

Stingless bees are globally important social corbiculate bees, fulfilling critical pollination roles in many ecosystems; however, their gut microbiota, especially fungal communities, are not well characterized to date. We collected 121 bee samples from two species, *Tetragonula carbonaria* and *Austroplebeia australis*, across a distance of 1,200 km of eastern Australia, and analysed their gut microbiomes. We found that the gut bacterial richness of *T. carbonaria* was influenced by geography (latitude and longitude) and positively correlated to an established fitness indicator in insects; namely, host forewing length/size that relates to flight capacity of stingless bees. We characterized the core microbiomes of the two bees and found that they consisted of the bacterial taxa *Snodgrassella, Lactobacillus*, Acetobacteraceae and *Bombella*, and the fungal taxa Didymellaceae, *Monocilium mucidum, Malassezia restricta* and *Aureobasidium pullulans*. Both host species identity and management (wild or managed) significantly influenced the gut microbial diversity and composition, and similarity between colonies declined as the geographical distance between them increased. This result was also supported by our co-existing network analyses. Overall, we have thoroughly analysed stingless bee gut microbiomes, and provided novel evidence that *T. carbonaria* bees with larger wings or from more southern populations have higher microbial diversity in their guts.

**Originality-Significance Statement:** Beneficial interactions between insects and their microbial symbionts are pivotal for their fitness. In this study, we analysed the gut microbiomes of two stingless bee species, *Tetragonula carbonaria* and *Austroplebeia australis*, that are widespread and important pollinators in Australia. We characterized their gut microbiomes and detected a significant positive correlation between gut bacterial richness and host forewing size for *T. carbonaria*; the first time that gut microbial diversity has been linked to a morphological trait in stingless bees. Furthermore, we found that host species’ identity, management type (wild or managed) and geography all significantly influenced bee gut microbial diversity and composition, and were able to describe both bacterial and fungal core microbial taxa. This study reveals novel understandings of stingless bee gut microbiomes and provides the basis for utilizing microbial strategies to maintain colony health.

## Introduction

Insect guts harbour many microorganisms across the three primary regions; foregut, midgut and hindgut (Chapman and Chapman, 1998). These microorganisms have various host functions that vary from aiding nutrient extraction from foods (Engel and Moran, 2013) to detoxification of harmful compounds (Ceja-Navarro et al., 2015) and protection against parasites and pathogens (Endt et al., 2010). Corbiculate bees in particular, are known to possess characteristic gut microbiomes. Honey bee (*Apis* spp.) guts, for example, consist of a core bacterial community including *Snodgrassella, Gilliamella, Lactobacillus* Firm-4 and -5 and *Bifidobacterium* (Koch and Schmid-Hempel, 2011; Kwong et al., 2017), which are obtained through floral visits (McFrederick et al., 2017; Liu et al., 2019) and from within the hive environment (e.g. through feeding on stored pollen). Increasing evidence shows that, similar to other insects, corbiculate bees have co-evolved with their gut microbes to form mutualistic relationships. The bees benefit from the microbiome primarily through defence against enemies, regulation of growth and development and contributing to hive pollen fermentation (Cano et al., 1994; Vásquez and Olofsson, 2009; Anderson et al., 2011; Koch and Schmid-Hempel, 2011; Engel et al., 2012; Menezes et al., 2015; Kwong et al., 2017). Conversely, changes to the gut microbiome composition of social bees, such as those caused by antibiotic exposure, can lead to dysregulated immune systems and reduced ecological fitness (Liu et al., 2019). Indeed, maintenance of a healthy gut microbiome is essential for the colony and individual’s homeostasis, nutrition and defence (Engel and Moran, 2013).

Among the corbiculate bees, stingless bees (tribe: Meliponini) comprise >500 species globally (Michener, 2013), of which 11 recognised species occur in Australia, under two genera: *Austroplebeia* and *Tetragonula* (Dollin and Dollin, 1997; Dollin et al., 2015). They are important pollinators of natural plants and crops (Heard, 1999; Hall et al., 2020), and can be harnessed by beekeepers either through rescuing colonies from felled trees, or propagation in man-made hives (Halcroft et al., 2013). The sale of stingless bee hives for crop pollination in Australia has increased over recent decades and, while natural populations exist across much of eastern Australia, most commercial colonies originate from a narrow region of southern Queensland (Chapman et al., 2018). Thus, the origin of the colony and whether it resides in a managed hive or is naturally occurring in a tree hollow, may affect how microbiomes are shaped e.g., by the internal conditions within hives (Heard 2016). *Austroplebeia* and *Tetragonula* stingless bees are similar in body size and colour and can occur sympatrically along the east coast of Australia. However, they belong to different phylogenetic clades, and *Austroplebeia* tends to occur further inland in somewhat drier habitats. Their behaviour also differs; for example, *T. carbonaria* is more active in flight and perhaps collects more resin and pollen than *A. australis* (Leonhardt et al., 2014). In contrast,

*A. australis* colonies are more likely to focus on collecting high-quality nectar (e.g., of high sugar concentrations) (Leonhardt et al., 2014). Such distinct behaviour, along with differences in available floral resources within their habitats can thus shape different gut microbiomes among stingless bee species (Vásquez et al., 2012). Previous studies have shown that bacterial communities within the two Australian stingless bee genera can change rapidly (Hall et al., 2021), and a novel clade of host-specific lactic acid bacteria (*Lactobacillus*) has been found (Leonhardt and Kaltenpoth, 2014; Hall et al., 2021). However, these studies used relatively few samples and to date there is limited comparison of gut microbial communities across species and geographic ranges. Additionally, like other animal gut microbiome studies, fungal communities in the guts of insects, including stingless bees, have received little attention (de Paula et al., 2021). Insect-associated fungi including moulds and yeasts can contribute to host nutrient provision (Menezes et al., 2015). For instance, some fungi are involved in wood and food digestion in underground ant and termite chambers (Zoberi & Grace, 1990). Similarly, the intracellular symbiotic fungi of beetles, *Symbiotaphrina* spp., can both aid in food digestion and detoxify a variety of plant materials (Dowd and Shen, 1990). Additionally, a Brazilian stingless bee *Scaptotrigona depilis* farms specific strains from the fungal genus *Monascus* (Ascomycota) as a food source for its larvae (Menezes et al., 2015). Despite the importance, fungal community composition and diversity, their mutualistic interactions with the host and key drivers of the microbiome assembly remain poorly understood.

The western (or European) honey bee (*Apis mellifera*) is one species for which a core bacterial microbiome has been determined (Kwong et al., 2017). The bacteria *Snodgrassella alvi, Gilliamella apicola, Lactobacillus sp*. and *Bifidobacterium* are ubiquitous and can be found in essentially every adult worker honey bee, with each bacterial species showing a specialized distribution in the gut (e.g., *S. alvi* and *G. apicola* dominate the ileum region of hindgut) (Moran, 2015). Each of the core species can contain multiple strains, with a high level of strain variation. Other social corbiculate bees, such as bumble bees and stingless bees, may contain distinct strains of similar bacterial species to those found in honey bees (Koch et al., 2013), but systematic studies of the core gut microbiome, in particular fungi and their interactions with other microbial species within the microbiome, are yet to be done.

One of the aims of this study was to test if correlations exist between stingless bee gut microbiome (bacteria and fungi) and morphological traits, such as flight capacity. Wing size in insects is an essential functional trait for flight performance (Wootton, 1992), foraging, dispersal and migration (Johansson et al., 2009). The wings of monarch butterflies (*Danaus plexippus*) from migratory populations, for example, tend to be larger than those from non-migratory populations, presumably because larger wings provide greater surface area for soaring flight and energy preservation (Dockx, 2007; Altizer and Davis, 2010; Davis and Holden, 2015). Insect wing length can predict flying capacity, and there is evidence that honey bees from colonies infected with the ectoparasite *Varroa destructor* had shorter wing lengths and flew shorter distances than healthy bees (Attisano et al., 2013; Blanken et al., 2015). Maximum flight distances of stingless bees were also highly correlated with wing size in six stingless bee species, suggesting that flight capacity is a function of their wing sizes, and thus, bees with larger wings may be able to fly further to forage on more diverse plant resources (Casey et al., 1985; Byrne et al., 1988; Araújo et al., 2004). However, to date no link has been made between stingless bee morphological traits, such as wing, tibia and body sizes, and gut microbial diversity.

Overall, despite the importance of stingless bees to pollination services, relatively little is known of their gut microbiome, nor how sorting and dispersal limitation function in shaping gut microbial communities; especially, how they are influenced by the host species, management type and geography. Here, we collected 121 samples of stingless bee foragers from two species; *Austroplebeia australis* and *Tetragonula carbonaria*, ranging 1,200 km along the east coast of Australia (Queensland and New South Wales), and investigated the composition of their gut microbiomes using 16S rRNA gene and fungal ITS amplicon sequencing. We tested the following hypotheses: (i) a core microbiome exists in the gut of stingless bee foragers; (ii) the gut microbiome structure is influenced by bee species, geographic location and by whether bees are wild or managed (i.e. cultivated in hives); and (iii) characteristics associated with flight and foraging capacities, such as wing length and size, positively correlate to host gut microbiome diversity.

## Material and methods

### Specimen collection, measurement and gut dissection

We collected 121 stingless bee samples from the two most common and widespread Australian stingless bee species, *T. carbonaria* and *A. australis*, across their distributional ranges in QLD and NSW, Australia (**Fig.1a**). The *T. carbonaria* samples (n=80) were collected between September 2018 and January 2020 across a range of 1,200 km in eastern Australia. They comprised 43 samples collected from managed hives and 37 samples collected from the wild (e.g. national parks). The *A. australis* samples (n= 41) had been collected previously (Halcroft et al., 2013), in 2009 from wild tree-living colonies across a 250 km range across their natural distribution (**Fig.1a**). Consequently, only *T. carbonaria* was used for analyses of the effect of management type (wild or managed) on gut microbial composition and diversity. During sampling, geographic coordinates (longitude and latitude) were recorded, and all live specimens were immediately preserved in 70% ethanol, and stored at −20 °C prior to gut dissection and morphological measurement. To minimize biases due to inter-individual variability, three individuals from each sample were used for microbiome analyses. The whole gut of each individual was dissected on a sterile Petri dish under a microscope using sterilized forceps, and the gut materials were then pooled and transferred to a 1.5 mL sterile centrifuge tube, and preserved at −20 °C prior to DNA extraction. We took the following morphometric measurements to infer bee flight performance and fitness (Wootton, 1992): forewing length, forewing area (forewing length × width), hind tibia length and total body length. Each trait was measured using a digital microscope (Leica EZ4W, Leica Microsystems, Buffalo Grove, IL) for 1∼3 bees per sample (2∼6 measurements of wings and hind tibia for each sample and an average of each trait was used as the final result of the sample).

**Fig.1.**
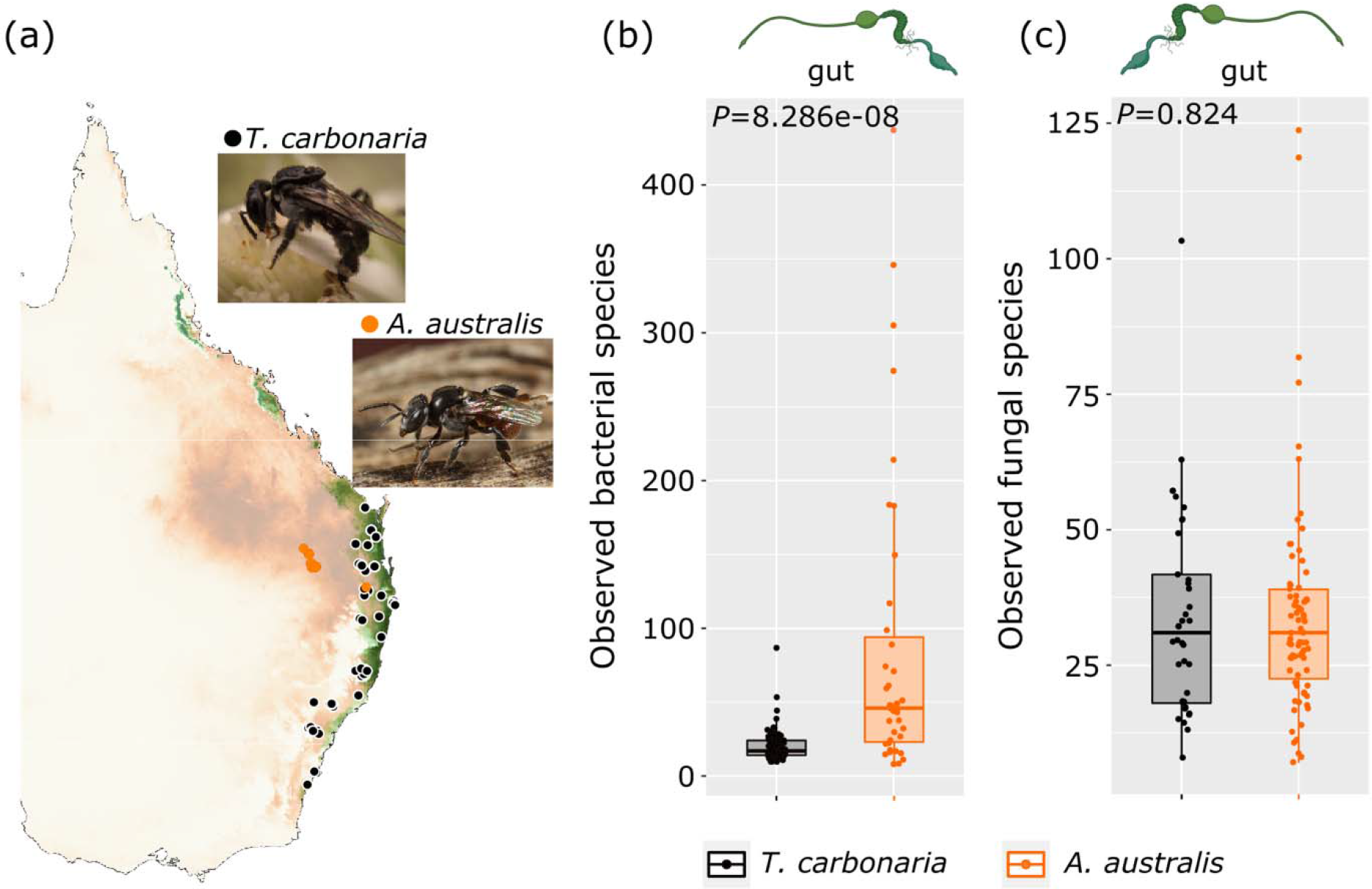
Sampling locations of stingless bee specimens across their known range in eastern Australia (a) and alpha diversity of their gut microbiomes: bacterial communities (b) and fungal communities (c). Solid black dots in (a) represent sampling locations for *T. carbonaria* while orange dots represent locations for *A. australis*. Green shaded areas represent the predicated range for *T. carbonaria*, while orange shading indicates the predicted east-coast range of *A. australis*.

### DNA extraction from gut material and library preparation for high throughput amplicon sequencing

For each sample, the gut material of three bees were combined and used for DNA extraction using the DNeasy Blood and Tissue Kit (Qiagen) as per the manufacturer’s recommendations. DNA samples were quality checked and quantified using a Nanodrop 2000 (Thermofisher Scientific, USA) and Qubit (Thermofisher Scientific, USA) respectively, before being stored at −20°C. Library preparation and bacterial and fungal amplicon sequencing were then carried out at the Next Generation Sequencing Facility (Western Sydney University, Australia). The 16S rRNA gene primers 341F (CCT ACG GGN GGC WGC AG) and 805R (GAC TAC HVG GGT ATC TAA TCC) (Herlemann et al., 2011) and the fungal ITS2 primers fITS7 (GTG ART CAT CGA ATC TTT G) and ITS4 (TCC GCT TAT TGA TAT GC) (Ihrmark et al., 2012; White et al., 1990) were used for the amplification and subsequent sequencing. All the samples were run on the MiSeq (Illumina) platform, generating 300 bp paired end reads.

### Bioinformatics and statistics

Sequencing files (FASTQ format) were processed using QIIME2 software and its plugins (version 2019.7; http://qiime2.org/) (Bolyen et al., 2019). Briefly, the sequencing quality was first assessed using FastQC (Andrews, 2010). Then, QIIME2 implementation of cutadapt v2019.7.0 was used for removal of primer sequences, and DADA2 v2019.7.0 (Callahan et al., 2016) was used for error-correction, quality filtering, chimera removal and constructing feature tables and final sequence files. Sequencing reads were truncated at 260 bp and 240 bp for forward and reverse reads, respectively, resulting in sequence quality Q>20. The amplicon sequence variants obtained were summarized and then assigned with taxonomic information using the q2-feature-classifier. For the bacterial data, a Naïve Bayes classifier pre-trained on the Greengenes 13_8 99% OTUs dataset was used to assign taxonomy to each representative sequence. Bee- and plant-associated mitochondria and chloroplast sequences were removed from the feature table to retain microbial features only. For the ITS fungal dataset, the classifier was trained to UNITE v8.0 database (99%) (UNITE Community, 2019) (DeSantis et al., 2006). The number of reads for the bacterial and fungal sequencing data was rarefied to 7,125 and 944 sequences, respectively, per sample by re-sampling the OTU table. The mean number of observed species (Obs), Chao1, Pielou_e, Simpson’s and Shannon diversity index values were calculated within QIIME2.

### Statistical analyses

R version 4.0.3 (2020-10-10) was used for analyses unless otherwise stated. Correlations between stingless bee traits and gut microbial alpha diversity were examined using linear models (Pearson) and visualised in R. The effect of stingless bee species and management types on gut microbial community composition and diversity were investigated using permutational multivariate analysis of variance (PERMANOVA, permutation=9999), and visualized with principal component analysis (PCA) using the Vegan package (v.2.5-6, (Oksanen et al., 2013). Fitting bee traits onto PCA ordination was performed using function *envfit* in Vegan (v.2.5-6). The ggplot2 package (version 3.3.3, (Ginestet, 2011)) was then used to produce the stacked graph at phylum level.

The phylogenetic tree for the most abundant OTUs was calculated in QIIME2, and annotated using Interactive Tree of Life (ITOL) (https://itol.embl.de). For core microbiome analysis and random forest test, we used an online microbiome analyses tool following recommended parameters (https://www.microbiomeanalyst.ca/MicrobiomeAnalyst/home.xhtml) (Chong et al., 2020). The determination of the core microbiome identifies those members that occur in >20% hosts at abundance of >0.1% within a defined host population, which is consistent with previously used methods (Bereded et al., 2020). For analysing gut microbial community variation over spatial gradients (latitude and longitude of each sample), we constructed geographic and environmental distance-decay relationships based on our spatially highly resolved set of samples (Soininen et al., 2007). This analysis reveals how the similarity in host microbiome composition between communities varies with geographic distance. The R package geosphere (1.5-10) (Hijmans, 2019) was used to calculate distance (km) between locations based on geographic coordinates for each sample. The *vegdist* function in the Vegan package (v.2.5-6) was used to calculate Bray-Curtis similarity (1-Bray-Curtis dissimilarity). Distance-based multivariate analysis for a linear model was then performed to investigate correlations between the Bray-Curtis similarity and geographic distance between stingless bee samples.

Lastly, the microbial co-existing network analysis was used to explore the interactions within the gut microbiomes. Co-existing network analysis reveals the potential interactions between microbial symbionts coexisting in the gut environment (Faust and Raes, 2012). Prior to microbial co-existence network calculation, OTUs present with <5 reads and in <3 samples in the normalized OTU tables were discarded to minimize spurious correlation that can be caused by rare taxa. Benjamini-Hochberg corrections (based on 100 bootstraps) were conducted to reduce the false discovery rate, and only strong (*R*>0.60) and robust (*P*<0.01) correlations were maintained. Correlations were calculated using the SparCC-based (Friedman and Alm, 2012) algorithm Fastspar (Watts et al., 2019), which was visualized and edited in Gephi (Bastian et al., 2009). Structural equation models (SEM) were used to evaluate the effects of morphometric traits and management approaches of bees on the bacterial richness in their gut, which was conducted using AMOS17.0 (AMOS IBM, USA). The maximum-likelihood estimation was fitted to the SEM model, and Chi-square and approximate root mean square error were calculated to examine model fit. Adequate model fits were determined according to a non-significant chi-square test (*P* > 0.05), high goodness fit index (GFI) (> 0.90), low Akaike value (AIC) and root square mean error of approximation (RMSEA) (< 0.05) as previously described (Delgado-Baquerizo et al., 2016).

## Results

### The alpha diversity of stingless bee gut microbial communities

The gut bacterial and fungal communities of the two stingless bee species across a substantial geographic range were characterized using high throughput 16S rRNA gene and ITS amplicon sequencing (**Fig.1a**). There was a significantly higher alpha diversity of the bacterial communities in *A. australis* than *T. carbonaria* (Obs *P*<0.0001 **Fig.1b**, Chao1 *P*<0.0001 and Shannon *P*=0.002, but not Simpson *P*=0.29). Similarly, alpha diversity of the gut fungal community of *A. australis* was significantly higher than that of *T. carbonaria* (Shannon *P*=0.017, Simpson *P*=0.033, Pielou_e *P*=0.0047, but not Obs *P*=0.82 **Fig.1c** or Chao1 *P*=0.83). For gut bacterial communities of *T. carbonaria*, wild bees had higher alpha diversity than managed bees (Chao1 *P*=0.04, but not for Obs *P*=0.063). Conversely, alpha diversity of the fungal community was higher in managed *T. carbonaria* bees than in wild bees (Chao1 *P*=0.003, observed species *P*=0.004).

### Core microbiome analyses of the stingless bee gut microbial communities

At the phylum level, gut bacterial communities of both stingless bee species were dominated by Proteobacteria, Firmicutes and Actinobacteria, along with less abundant Bacteroidetes, Verrucomicrobia, Tenericutes, Acidobacteria, Gemmatimonadetes and other unidentified taxa (**Fig.S1a**). The fungal community was dominated by Ascomycota and Basidiomycota, with Chytridiomycota, Mucoromycota and some unidentified fungal phyla also common (**Fig.S1b**). To obtain an overall picture, we combined samples of the two species and analysed their phylogenetic traits and dominant phylotypes using the top 255 (in relative abundance) bacterial sequences (**Fig.2a**). We found that, at the OTU level, the stingless bee gut microbiome was largely dominated by five OTUs, namely g_*Snodgrassella* (16.2%), f_Acetobacteraceae (10.3%), f_Acetobacteraceae (9.8%), g_*Lactobacillus* (8.6%) and g_*Lactobacillus* (6.1%) (**Fig.2a**). The stingless bee gut microbiome composition appears to be highly variable within host species and colonies. When the presence threshold was set at 20% for defining the core microbiome, we found seven core bacterial OTUs, including: g_*Snodgrassella*, f_Acetobacteraceae, g_*Lactobacillus*, f_Acetobacteraceae, g_*Lactobacillus*, g_*Bombella* and s_*Lactobacillus* sp. Thmro15 (**Fig.2b**), and six fungal OTUs: f_Didymellaceae, s_*Monocilium mucidum*, s_*Malassezia restricta*, s_*Aureobasidium pullulans*, s_*Alternaria alternata* and s_*Malassezia globosa* (**Fig.2c**). The core gut microbiome of these Australian stingless bees consists of a distinct group of bacterial taxa from that observed in honey bees and bumble bees (**Table 1**). However, there are currently no available data on core fungal taxa of other social corbiculate bees to compare. Fungal guild analyses identified the core fungal species in stingless bees as both litter/soil saprotrophic and also invertebrate-associated taxa (**Table 1**).

**Fig.2.**
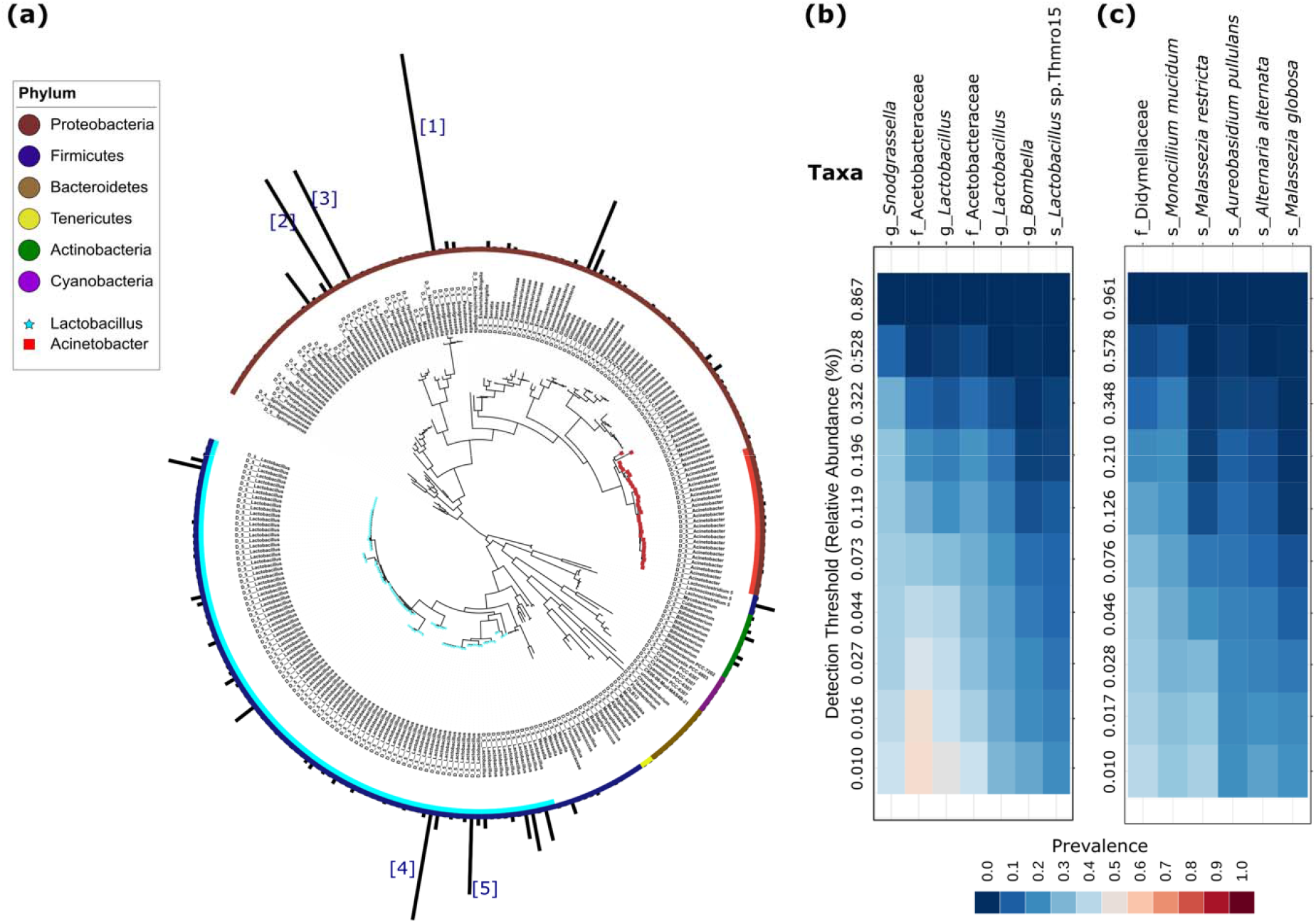
The phylogenetic tree of the stingless bee gut microbiome and core microbiome composition. Phylogenetics and the abundance of the top 255 OTUs of the gut bacterial taxa (a). The colour of each of the phyla was marked in different colours as indicated in the top label. The dominant bacterial genera of *Lactobacillus* and *Acinetobacter* are marked by star and square, respectively. Numbers in brackets represent [1] g_*Snodgrassella* (comprising 16.2% of total taxa), [2] f_Acetobacteraceae (10.3%), [3] f_Acetobacteraceae (9.8%), [4] g_*Lactobacillus* (8.6%) and [5] g_*Lactobacillus* (6.1%). Core microbial species (prevalence >20%) in the gut are shown for bacterial species (b) and fungal species (c), respectively.

**Table 1.** Comparison of core gut microbiome in the three main eusocial bee groups.

| Honey bee (*Apis* spp.)<br>(Kwong et al., 2017) |  | Bumble bee (*Bombus* spp.)<br>(Kwong et al., 2017) |  | Stingless bee (*Tetragonula* and *Austroplebeia* spp.)<br>(this study) |  |  |
| --- | --- | --- | --- | --- | --- | --- |
| Bacteria | Fungi | Bacteria | Fungi | Bacteria | Fungi | Fungal guild |
| *Snodgrassella* | Not<br>well<br>characterized | *Snodgrassella* | Not<br>well<br>characterized | *Snodgrassella* | Didymellaceae | NA |
| *Gilliamella* |  | *Gilliamella* |  | Acetobacteraceae | *Monocillium mucidum* | Litter saprotroph/<br>invertebrate-associated |
| *Lactobacillus*<br>Firm-4 |  | *Lactobacillus*<br>Firm-4 |  | *Lactobacillus* | *Malassezia restrita* | soil_saprotroph |
| *Lactobacillus*<br>Firm-5 |  | *Lactobacillus*<br>Firm-5 |  | *Lactobacillus* sp.<br>Thmro15 | *Aureobasium pullulans* | Sooty mould |
| *Bifidobacterium* |  | *Bifidobacterium* |  | *Bombella* | *Alternaria alternata* | Plant pathogen |
| *Bartonella apis* |  | *Bartonella apis* |  |  | *Malassezia globosa* | Soil saprotroph |
| *Frischella* |  | *Frischella* |  |  |  |  |
|  |  | *Bombiscardovia* |  |  |  |  |
|  |  | *Schmidhempelia* |  |  |  |  |

### Co-existing network analyses of the stingless bee gut microbiomes

Bacterial and fungal taxa, and geographic coordinates (latitude and longitude) of samples were integrated for co-existing network analyses to detect potential microbe-microbe interactions and predict the involvement of geography in the microbial network (**Fig.3**). From 121 samples and 2,799 OTUs, correlation network analyses identified 81 strong correlations (*R*> 0.6, *P* < 0.01) for managed *T. carbonaria* (**Fig.3a**), 95 for wild *T. carbonaria* (**Fig.3b**) and 574 for wild *A. australis* (**Fig.3c, Table S1**). The three networks (*T. carbonaria* managed and wild and *A. australis* wild) were formed by six distinct major modules, and each module represents a group of OTUs that are closely connected with each other but less linked with other OTUs in the network. The average degree of correlation for each of the three networks ranged from 2.76 to 8.43, with *A. australis* having the largest degree (8.43) followed by *T. carbonaria* wild (3.84) and *T. carbonaria* managed (2.76) (**Table S1**), suggesting that wild *A. australis* may have the most extensive microbial interactions of the three groups.

**Fig.3.**
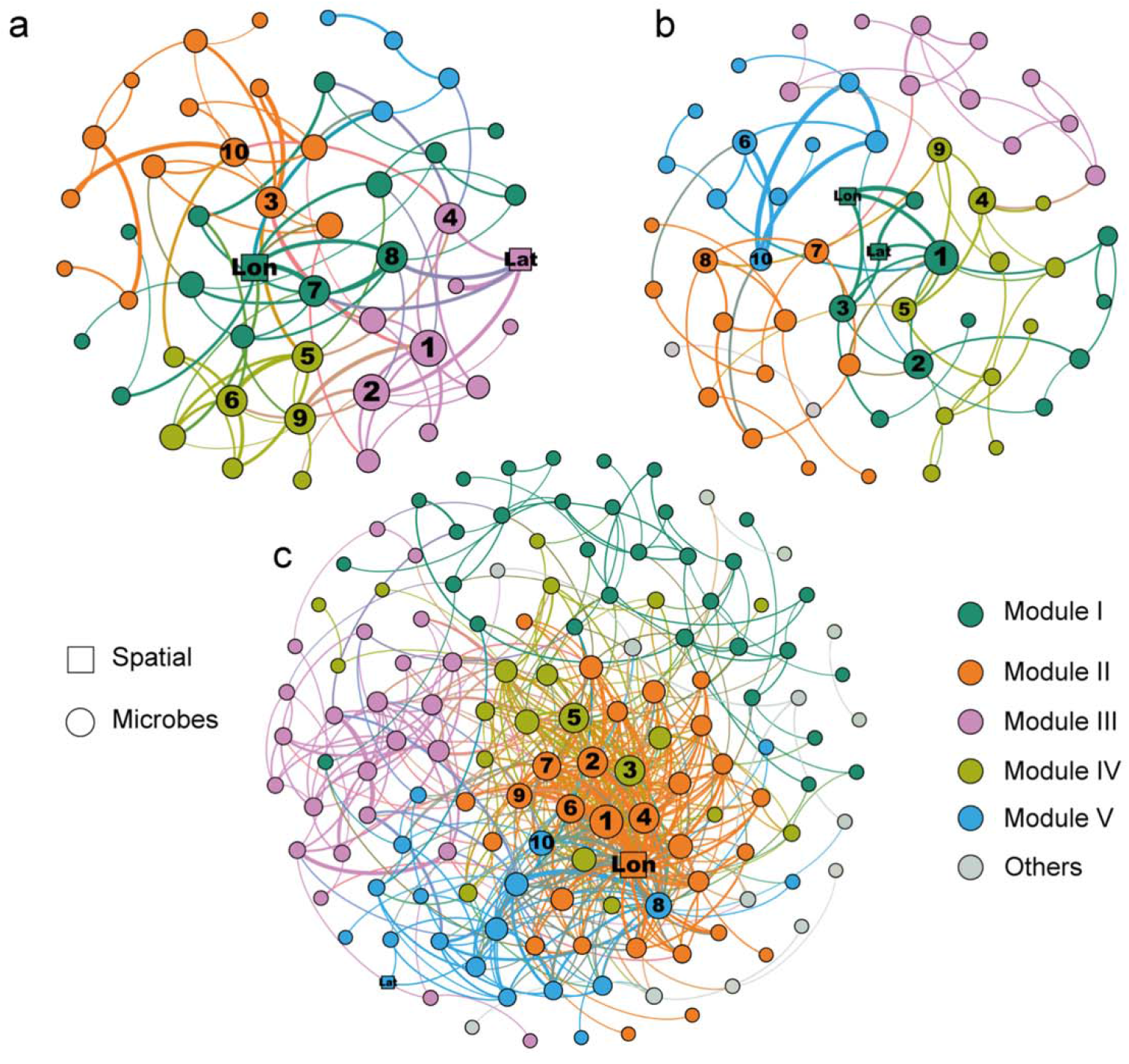
Network analyses of stingless bee gut microbiomes. Network analysis reveals the co-existing patterns among bacterial and fungal taxa and geographic coordinate. Connections represent a strong (Spearman’s correlation coefficient *R*> 0.6) and significant (*P*< 0.01) correlations. The size of each node is proportional to the number of connections. Networks shown are for managed *T. carbonaria* (a), wild *T. carbonaria* (b) and wild *A. australis* (c) colonies. Numbers [1]-[10] within each network represent the top 10 most connected bacterial/fungal taxa to other OTUs in the gut microbiome: *T. carbonaria* managed: [1] g_*Bombella*, [2] g_*Zymobacter*, [3] s_*Neophaeomoniella niveniae* (fungi), [4] s_*Alternaria alternata* (fungi), [5] g_*Lactobacillus*, [6] g_*Weissella*, [7] s_*Pseudosydowia eucalypti* (fungi), [8] g_*Hormonema* (fungi), [9] s_*Filobasidium oeirense* (fungi) and [10] g_*Gilliamella*; *T. carbonaria* wild: [1] s_*T. carbonaria*, [2] g_*Rosenbergiella*, [3] g_*Lactobacillus*, [4] g_*Acinetobacter*, [5] g_*Snodgrassella*, [6] s_*Lactobacillus* sp. Thmro15, [7] s_*Pseudosydowia eucalypti* (fungi), [8] s_*Quambalaria cyanescens* (fungi), [9] g_*Zymobacter*, and [10] longitude (geographic). *A. australis*: [1] f_Rhodobacteraceae, [2] f_Burkholderiaceae, [3] g_*Algoriphagus*, [4] g_*Hydrogenophaga*, [5] longitude (geographic), [6] s_*Terrimonas* sp. 16-45A, [7] g_*Cyanobium* PCC-6307, [8] g_*Porphyrobacter*, [9] g_*Snodgrassella* and [10] g_*Alishewanella*. Lon: longitude, Lat: latitude.

The 10 microbial taxa with the most connections with other microbes were highlighted in each network structure (**Fig.3a,b,c**). Such microbial taxa within the managed *T. carbonaria* network included g_*Bombella*, g_*Zymobacter*, s_*Neophaeomoniella niveniae* (fungi), s_*Alternaria alternata* (fungi) and g_*Lactobacillus* (**Fig.3a**). The most connected microbial taxa in the network of wild *T. carbonaria* included s_*Bifidobacterium bombi*, g_*Rosenbergiella*, g_*Lactobacillus*, g_*Acinetobacter* and g_*Snodgrassella* (**Fig.3b**). For wild *A. australis*, these included f_Rhodobacteraceae, f_Burkholderiaceae, g_*Algoriphagus*, g_*Hydrogenophaga*, and s_*Terrimonas* sp. 16-45A (**Fig.3c**). We also found a few microbes with close connections to latitude and longitude, indicating a geographic influence in the formation of the gut microbiome network. Among these, there were three taxa with connections to latitude and longitude, for managed *T. carbonaria* (**Fig.3a**), 5 and 7 connections respectively for the wild *T. carbonaria* (**Fig.3b**). For *A. australis*, there were 2 taxa with connections to latitude and 33 to longitude (**Fig.3c**).

### Factors influencing stingless bee gut microbial community composition (beta-diversity)

Bacterial and fungal community composition were influenced by both host species identity (Bacterial: *R*^2^=0.19, *P*=0.0001, fungal: *R*^2^=0.03, *P*=0.0001) and management type (managed/wild, assessed in *T. carbonaria* only) (Bacterial: *R*^2^=0.025, *P*=0.023, fungal: *R*^2^=0.064, *P*=0.0001) (**Figs.4a,b**). When both bee species were considered together, the gut bacterial community composition was closely associated with host forewing length (*R*^2^=0.2047, *P*=0.0001), latitude (*R*^2^=0.1420, *P*=0.0004) and longitude (*R*^2^=0.5051, *P*=0.0001) (**Fig.4a**). However, none of these factors were significant when the two species were analysed independently. For the fungal community, host forewing and tibia lengths and longitude were associated with the microbial community composition (*R*^2^=0.18, *R*^2^=0.40, *R*^2^=0.37 respectively, and *P*=0.0001 in all cases) (**Fig.4b**). When examining each species separately, body length was associated with the fungal community composition of *A. australis* (*R*^2^=0.18, *P*=0.034), while longitude was associated with the fungal community composition of *T. carbonaria* (*R*^2^=0.16, *P*=0.004).

**Fig.4.**
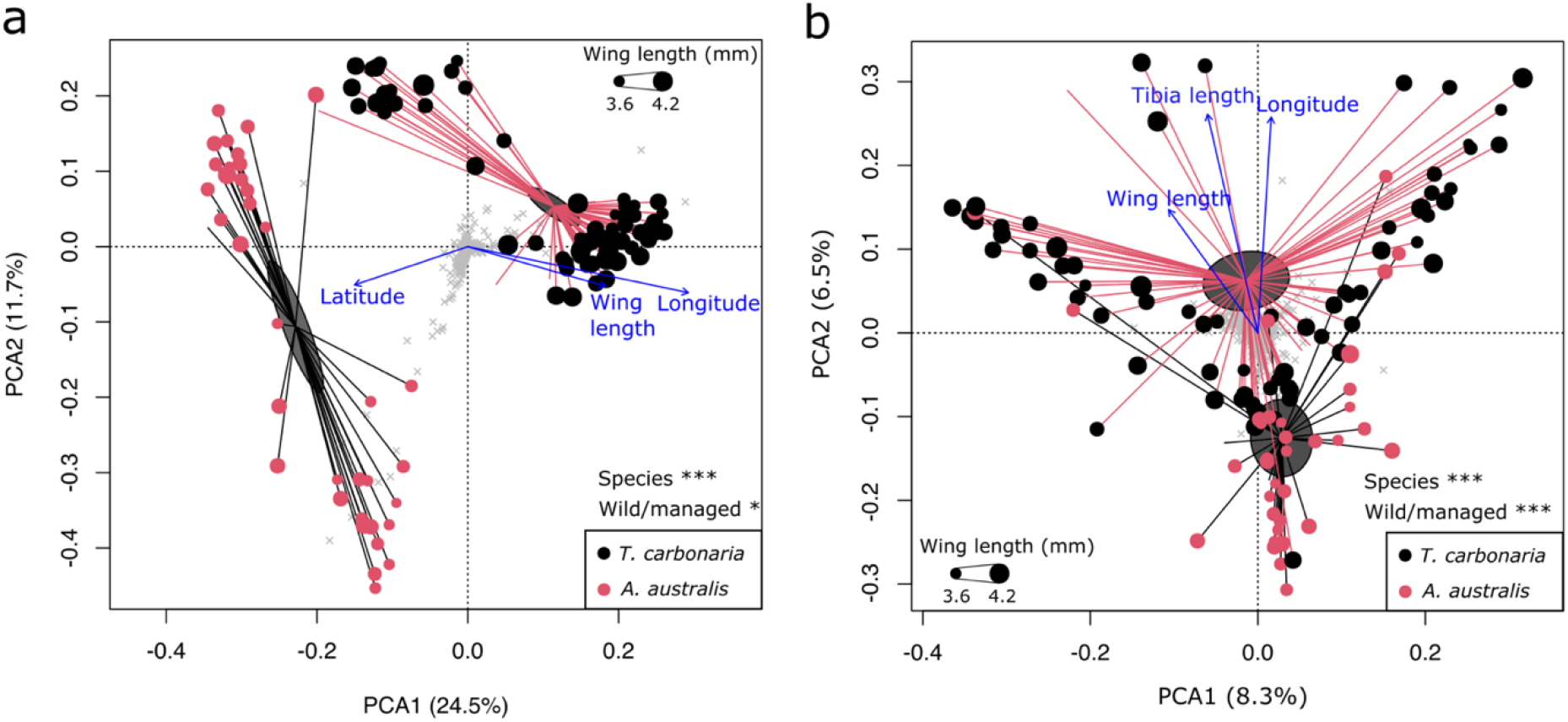
Principal component analyses (PCA) of the gut microbial community composition associated with *T. carbonaria* and *A. australis*: bacterial communities (a), and fungal communities (b). Ellipses show the standard error of the mean (SE). Blue arrows represent the direction of environmental gradients, with length proportional to strength of the correlation.

We then identified key microbial taxa distinguishing the gut microbiomes of the two species using random forest tests (**Fig.S2**). Error rates for the bacterial and fungal communities achieved stability when tree numbers for prediction reached 110 and 140 for bacterial and fungal communities, respectively (**Fig.S2a,b**). The top ten bacterial OTUs that distinguished between the two gut microbial communities were revealed; one belonging to g_*Gilliamella*, six g_*Lactobacillus*, one g_*Mycobacterium* and one s_uncultured bacterium were associated with *A. australis*, whereas one OTU belonging to f_Acetobacteraceae was associated with *T. carbonaria*. For the fungal community, one OTU of s_*Zygosaccharomyces rouxii*, one f_Didymellaceae and seven unidentified fungal OTUs were associated with *A. australis*, whereas one OTU belonging to s_*Aureobasidium pulluans* was associated with *T. carbonaria*.

We also used Bray-Curtis similarity index to determine the influence of geographic distance (km) between samples on gut bacterial and fungal community composition (**Fig.S3**). A significant linear decrease in composition similarity with increasing geographic distance was seen between samples for *T. carbonaria* gut bacterial communities (*P*<0.0001). In addition, there was a similarity decrease with increasing geographic distance for fungal communities for both *T. carbonaria* (*P*<0.0001) and *A. australis* (*P*=0.002) (**Fig.S3a,c,d**). These results show that the microbiome composition varies more between samples from geographically distant locations. However, the adjusted R^2^ for each correlation was small (0.005∼0.01), indicating only a fairly weak link. No significant correlation was seen for gut bacterial communities in *A. australis* (**Fig.S3b**). It is worth noting that the *A. australis* samples covered a relatively smaller area, and so the power to detect small changes may have been reduced compared to *T. carbonaria*.

### Correlation of stingless bee gut microbial richness with host morphological traits and geography

Linear models showed that forewing length and area, hind tibia length and body length were all positively correlated (e.g. *R*_wing length–wing area_= 0.86; *R*_wing length–body length_=0.42; *R*_wing length–tibia length_=0.78; *N*=110; *P*<0.0001 in all cases) (**Fig.S4a,b**). Additionally, hind tibia (*P*<0.0001) and forewing length (*P*<0.0001) differed between the two bee species, but full body length and forewing area did not. Interestingly, gut bacterial richness (Obs) in *T. carbonaria* showed a significant positive correlation with host forewing length (*R*=0.44, *N*=71, *P*=0.001) and forewing area (*R*=0.32, *N*=71, *P*=0.008) (**Fig.5a**), while full body length and hind tibia length showed no correlation with gut bacterial richness or any other alpha diversity value. The forewing area/body length ratio, which is believed to determine flight capacity, also correlated with gut bacterial richness in *T. carbonaria* (*R*=0.26, *P*=0.033). For *A. australis*, however, no correlation was seen between alpha diversity of gut microbiomes and host traits, such as forewing length (bacterial *R*=−0.31, *P*=0.058; fungal *R*=0.12, *P*=0.49) or forewing area (*R*=−0.16, *P*=0.33; *R*=0.07, *P*=0.48).

**Fig.5.**
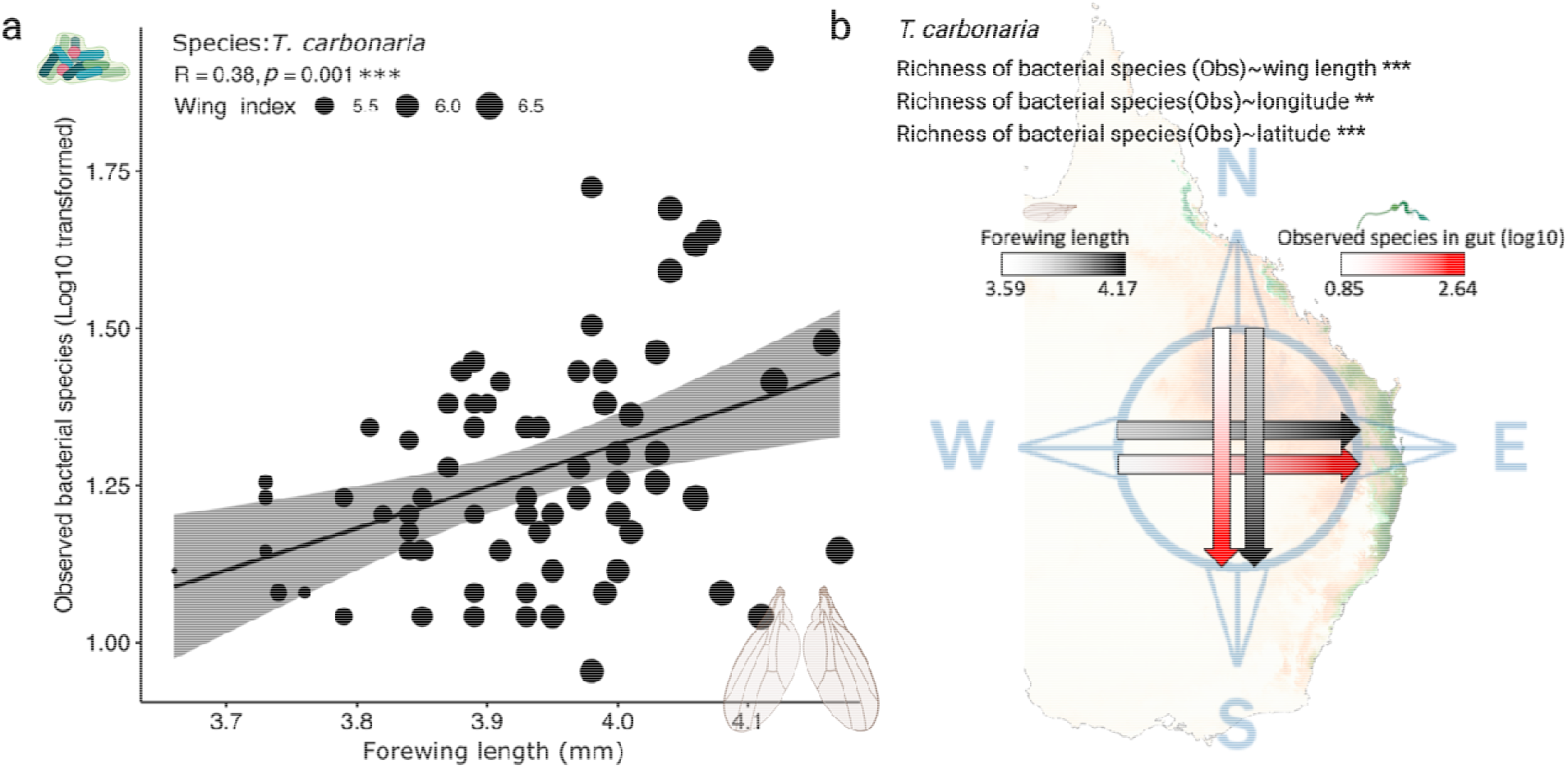
Linear correlation between gut bacterial species richness in *T. carbonaria* with host forewing length (a) and geographical location (b). Shown are regression (black line), 95% confidence intervals (shading) and data points (black dots). Wing length analyses shown at log10 scale.

We then analysed the relationship between geography and host gut microbial richness. Importantly, for *T. carbonaria*, longitude (*R*=−0.36, *P*=0.001) and latitude (*R*=−0.43, *P*<0.0001) negatively correlated with the gut bacterial species richness, but not with fungal species richness (**Fig.5b**). Longitude and latitude both influenced forewing length (Lon *R*=−0.38, *P*=0.0008; Lat *R*=−0.44, *P*=0.0001) and forewing area (Lon *R*=−0.38, *P*=0.0008; Lat *R*=−0.47, *P*<0.0001) (**Fig.5b**). For *A. australis*, neither latitude nor longitude influenced host morphological traits, which is not surprising given the sampling range for this species was considerably smaller than for *T. carbonaria*. Using SEM analysis that considered morphological traits, geographical factors and management effects together, we found a dominant effect of forewing length on gut bacterial richness, while longitude and latitude effects on gut microbial richness were small (**Fig.S5**).

## Discussion

We characterized the whole gut microbiome of two stingless bee species, *T. carbonaria* and *A. australis*, from 121 samples spanning a large geographic range (1,200 km) in eastern Australia. We tested whether gut microbiomes are linked to geographic location (longitude and latitude), host species, management type (wild vs managed) and host traits. For *T. carbonaria*, we observed a positive correlation between both forewing length and area with host gut bacterial richness; samples collected in the south had both longer wings and higher bacterial richness than those from the north of their range. For both species, microbiomes consistently became more distinct from one another in their composition with increasing geographic distance between samples. While this effect was relatively weak, it suggests a role of geographical factors in shaping stingless bee gut microbiomes. We analysed key microbial taxa using both core microbiome (abundance based) analyses and microbial co-existing network analyses, which revealed key bacterial groups that show differences from core microbes present in honey bee and bumble bee guts. This study is the first comprehensive analysis of stingless bee gut microbiomes. Combining analyses of both the bacterial and fungal communities with host traits and geography, we provide evidence of a link between the gut microbiome and host morphological traits in stingless bees.

### Stingless bee gut microbial diversity correlates with host forewing size and geographic location

We observed a positive linear correlation between gut bacterial species richness and host forewing length and size for *T. carbonaria*. Insect wing sizes are closely related to their flight capacity (DeVries et al., 2010); longer wings favour wider variation in speed and increase capacity for longer flight duration as well as perhaps also saving energy (DeVries et al., 2010). These factors potentially increase the capacity of bees to collect diverse floral resources. Furthermore, insects with larger wings are also more successful in host-seeking and their location of oviposition sites (Berwaerts et al., 2002; Davis and Holden, 2015). Therefore, larger wing size is likely to confer a fitness advantage. We did not, however, see any correlation between gut bacterial/fungal richness with body/tibia size, so the correlation with wing size is not an indirect consequence of a general correlation with body size. This study therefore provides solid evidence of a correlation between wing traits with the host gut microbiome. Additional research is now needed to determine whether increasing wing size results in an increase to the diversity of the gut microbiome, or *vice versa*. Furthermore, it is also important to consider the complexities of the tripartite interactions between wing size, gut microbial diversity and geography, especially considering geography, as measured here by latitude and longitude, actually involves a myriad of other environmental factors. For example, much of Queensland experiences a tropical/subtropical environment, whereas further south e.g., Western Sydney, experiences far greater temperature extremes. Thus, the physiological requirements of the bees, and the floral resources available to them will vary considerably.

### Bacterial communities

In this study, Proteobacteria and Firmicutes were found to be the dominant bacterial phyla in the stingless bee gut, followed by Actinobacteria; a pattern also observed in honey bees and bumble bees (Kakumanu et al., 2016; Wang et al., 2019). Such microbial similarity among species supports a strong host selection of the microbial environment by eusocial bees. The core bacterial phylotype, *Lactobacillus*, has important functions in the host, such as protection against pathogens, food digestion and pollen fermentation, as has been demonstrated for honey bees in previous studies (Engel and Moran, 2013; Liu et al., 2019). They are also common in the gut system of bumblebees worldwide (Kwong and Moran, 2016), suggesting that mutualisms with *Lactobacillus* exist throughout the eusocial bees across different geographic regions. Our random forest model showed that *Lactobacillus* spp. are the main indicator taxon distinguishing the stingless bee gut microbiomes of the two species, with six of the ten key distinguishing OTUs belonging to this genus. This suggests a great variance in phylogeny and/or abundance of the *Lactobacillus* genus at species/strain levels between the two species. How this diversity of *Lactobacillus* relates to biological differences and host function is yet to be studied.

*Snodgrassella spp*., another core bacterial genus in the stingless bee gut microbiome, also features in the core microbiome of both honey bees and bumble bees (Koch and Schmid-Hempel, 2011). *Snodgrassella* spp. are saccharolytic fermenters and interestingly, have been implicated in the protection of bumble bees against *Crithidia bombi* (Koch and Schmid-Hempel, 2011). Furthermore, laboratory studies indicated that glyphosate (the primary herbicide worldwide) can perturb the strain abundance of core gut *Snodgrassella alvi* in honey bees, which led to higher rates of mortality when glyphosate-treated bees were exposed to the opportunistic pathogen *Serratia marcescens*, highlighting the importance of this bacterium in the maintenance of host health. As with *Lactobacillus*, the relevance of the species/strain diversity of *Snodgrassella* spp. in the *stingless bee* gut microbiome is not yet understood, but may correspond to differences in host metabolic capabilities. Interestingly, a recent study surveyed gut microbiomes of Brazilian stingless bees by sampling multiple species within the genus *Melipona*, and showed that stingless bees can lose the core symbioses of *Snodgrassella* (Cerqueira et al., 2021). This suggests that strong ecological shifts or functional replacements in the stingless bee gut microbiome can occur.

Stingless bee guts also contained bacterial genera such as *Pantoea, Sphingomonas* and *Stenotrophomonas*, which commonly colonize all parts of flowering plants and are also found in hive environments (Liu et al., 2017; Liu et al., 2020). *Snodgrassella, Gilliamella* spp. (Graystock et al., 2017), *Saccharibacter* spp. (McFrederick et al., 2012), *Massilia* spp. (Graystock et al., 2017) and *Acinetobacter* spp. (Graystock et al., 2017) are commonly found on flowers (Liu et al., 2019), suggesting floral visits are key to the microbial acquisition by stingless bees.

The gut bacterial species richness of *A. australis* was significantly higher than that of *T. cabonaria*. Such microbial difference in gut may relate to the distinct foraging behaviour of the two species. *T. carbonaria* evidently collects more protein enriched food (e.g. pollen) than *A. australis* which likely focuses on high-quality nectar (carbohydrate enriched) (Leonhardt et al., 2014). This higher level of carbohydrate foraging may be linked with the higher bacterial richness seen in *A. australis* gut. Hive managed bees seem likely to possess less gut bacterial diversity than wild bees, which may indicate a less diverse food composition. A previous study also found that gut bacterial diversity of fruit fly (*Bactrocera tryoni*) larvae was significantly lower in laboratory populations compared with field populations (Deutscher et al., 2018). We also observed that stingless bee gut microbiomes vary greatly among samples. This aligns with a previous study on whole-body bacterial and fungal communities of managed *T. carbonaria* (Hall et al., 2021). Rapid temporal changes of the bee microbiome composition observed in that study may also, to some extent, explain the high variability of microbiome composition we saw across a geographic gradient. For example, Hall et al. (2021) saw dramatic increases in the relative abundances of *Bombella* and *Zymobacter* and almost complete depletion of *Snodgrassella* when colonies were moved from a florally resource-rich site to a resource-poor site. Other studies have also shown bacterial community diversity to vary considerably between bee colonies (Koch and Schmid-Hempel, 2011; Leonhardt and Kaltenpoth, 2014). All the above findings suggest that stingless bees are prone to compositional shifts, putatively influenced by food resources, physiological status, origin of the colony and climate at different geographic locations.

### Fungal communities

The dominant fungal phyla observed, Ascomycota and Basidiomycota, usually fulfil a decomposing role in most land-based ecosystems, by breaking down organic materials such as large molecules of cellulose or lignin, and in doing so play important roles in carbon and nitrogen cycling (Dighton, 2016). It is logical then, to assume that these gut colonisers can assist the animal host with food digestion (de Paula et al., 2021). We identified six core fungal taxa in *T. carbonaria* and *A. australis*. Interestingly, two of these, *Malassezia restricta* and *M. globosa*, are also among the most abundant fungal species in the human gut (as indicated by their large presence in faecal samples and intestinal mucosa), and have been identified in association with gut diseases including colorectal cancer (Coker et al., 2019). This may make stingless bees a suitable animal model for investigation of the role of these fungi in human gut health. Another core fungal genus, *Monocillium* spp. has previously been isolated from soil, dead leaves and wood and some species (e.g., *M. curvisetosum*) originate from aphids. There is evidence that *Monocillium* spp. are able to antagonise a plant parasitic nematode by colonising their cysts (Ashrafi et al., 2017). *Aureobasidium pullulans* was previously isolated from honey and honey bee guts (Good et al., 2014). The detected *Alternaria alternata* can be an opportunistic fungal pathogen on plants causing leaf spots, rots and blights (Tsuge et al., 2013); however, its function in the bee gut and whether it may be vectored between plants by bees is currently unknown. Further investigations are needed to determine how gut-colonising fungi interact with co-existing bacteria, and the implications for host nutrition and fitness. Nevertheless, the present study is of considerable value and interest in pollinator biology, providing insight into the gut-specific core fungi, and will enable future investigations into the functional role of the core fungal community associated with stingless bees.

### Distance-decay relationship between stingless bee gut microbiomes and geographic distance of bee sampling

Our data showed that microbial biogeographic patterns (a distance-decay relationship) could be applied to stingless bee gut microbiomes on a geographic scale of 250∼1,200 km. While we predicted decreasing community similarity over distance due to dispersal limitation and local adaptation of stingless bee microbiomes (Soininen et al., 2007; Nemergut et al., 2013), evidence for such a relationship in bee microbiomes was previously lacking. An analysis of relative abundances of *Snodgrassella* and *Gilliamella* across *Bombus* and *Apis* hosts found poor correlation with geography (Koch et al., 2013). Previous studies found limited effects of geographic location on microbiota composition probably due to small sample sizes and/or geographic distances (Kwong et al., 2017). However, the bacterial and fungal distance-decay relationship detected in our study, although significant, explained a small amount (R^2^=0.005∼0.01) of observed variation, perhaps smaller than those typically observed for plants and other animals. The number of samples, geographic area covered and sequencing depth all could affect the differences, which highlights the need to couple high-throughput sequencing methods with wide geographical coverage.

### Microbial interactions in stingless bee gut as estimated based on co-existing network analyses

The microbial network structure and complexity differed between the two stingless bee species, which can reflect distinct microbial interactions in their gut environments. However, whether these differences result in altered gut functions is not clear. The top five well-connected nodes within the network structures in managed and wild *T. carbonaria* were dramatically different from those identified in *A. australis*. The *T. carbonaria*-associated bacterial taxa, *Bombella, Lactobacillus, Acinetobacter* and *Snodgrassella*, have functional effects on nutrient acquisition and defence in honey bees and bumble bees (Kwong and Moran, 2016). In contrast, the potential role of the top five nodes - f_Rhodobacteraceae, f_Burkholderiaceae, g_*Algoriphagus*, g_*Hydrogenophaga* and s_*Terrimonas* sp. - in microbial networks of *A. australis* are not well characterized.

Our sampling strategy covered large geographic gradients in habitat conditions (*T. carbonaria* in particular), allowing for detection of robust co-existing patterns between microbes and their geographic locations. We showed that the five modules in the networks correlated with spatial variables, with each coordinate variable (latitude/longitude) being correlated with at least one module, indicating a potential role of geography in the formation of the network structure of the gut microbiome. To some extent, this supports our observation of a significant distance decay of the gut microbiome similarity for stingless bees. Whilst it does not necessarily imply cause, co-existing network analyses provide evidence for the role of location in governing stingless bee microbiome assembly (Faust and Raes, 2012).

## Conclusions

We characterized the gut microbiomes of two stingless bee species from different genera across 1,200 km thereby spanning large parts of their geographic ranges in eastern Australia. We observed large variance in the microbiome structure between species, which was also influenced by geographic location. Their gut microbial richness correlated to key host morphological traits, namely wing length and area. We have also provided novel evidence of the core fungal and bacterial species present in stingless bees, which appear to be distinct from honey bees, bumble bees and other insects. The link between gut microbiomes and bee fitness traits in *T. carbonaria* provide a novel framework to test functional interactions between bees and gut microbiome, and to develop future solutions to conserve and manage bees for crop pollination.

## Data availability

The raw sequencing data have been deposited in the NCBI Sequence Read Archive (SRA) under Bioproject code PRJNA698658. Datasets generated in this study are available at 10.6084/m9.figshare.14559801.

## Author contributions

HL, BKS, JC, RS, MR and LB conceived the idea; Mark H, Megan H, SN and HL collected wild and managed bee samples along eastern Australia; HL conducted bee gut dissections, bee measurement and DNA extraction, and analysed the sequencing data; Mark H drew Fig.1a and JW conducted the microbial co-existing network and SEM analyses; HL led writing of the manuscript, and all authors contributed to manuscript edits and approved final version for submission.

## Acknowledgements

This work was supported by the Hort Frontiers Pollination Fund [PH15001] (Healthy bee populations for sustainable pollination in horticulture) and host microbiome and multitrohic interaction fund (Australian Research Council (DP190103714; DP210102081)). We thank Dr Jasmine Grinyer for contributing a stingless bee sample from the Blue Mountains. Megan also provided records collected by the late Allan Biel, who recovered hundreds of stingless bee colonies, particularly around the Tara region of QLD. We thank him for his passion and commitment to Australian native bees.

