## Supplementary material for "Gut microbial diversity in stingless bees is linked to host wing size and is influenced by geography": na

**Table S1** Topology of microbial co-existing network analyses.

| | *T. carbonaria* (managed) | *T. carbonaria* (wild) | *A. australis* (wild) |
|---|---|---|---|
| Nodes | 58 | 49 | 139 |
| Edges | 80 | 94 | 573 |
| Average degree | 2.76 | 3.84 | 8.43 |
| Positive rates (0~1) | 0.71 | 0.51 | 0.62 |
| Modularity* | 0.61 | 0.47 | 0.36 |

*Modularity: a network property depicting the extent to which nodes are clustered with each other.

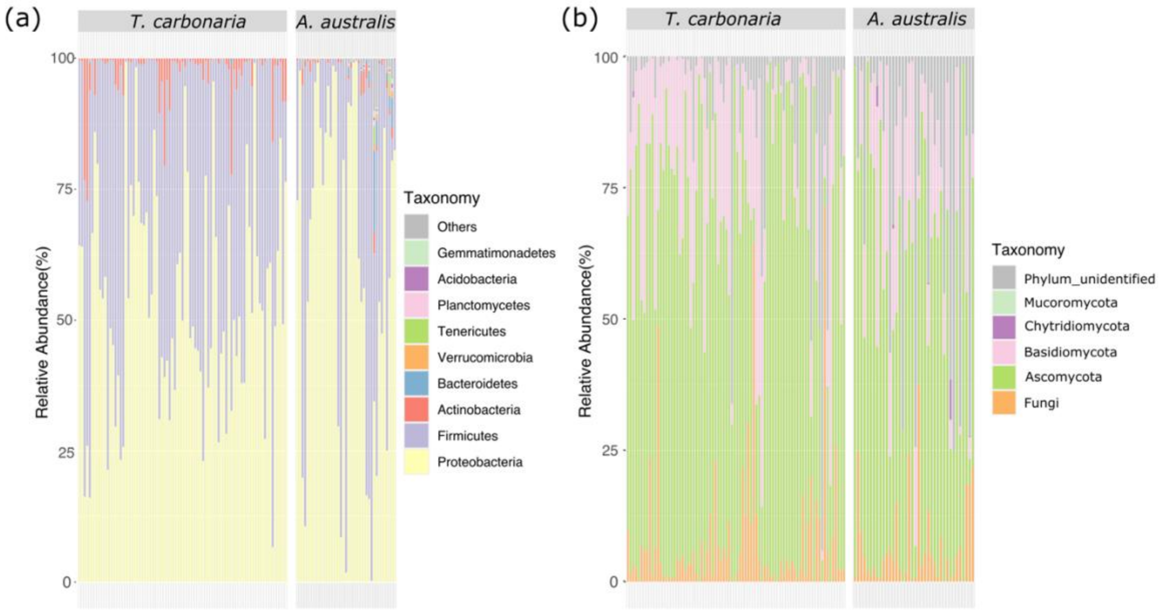


**Fig.S1 Stacked bar chart summarizing taxonomic composition of the stingless bee microbiome at the phylum level**: Bacterial community (a) and Fungal community (b). Each column represents a sampling location.

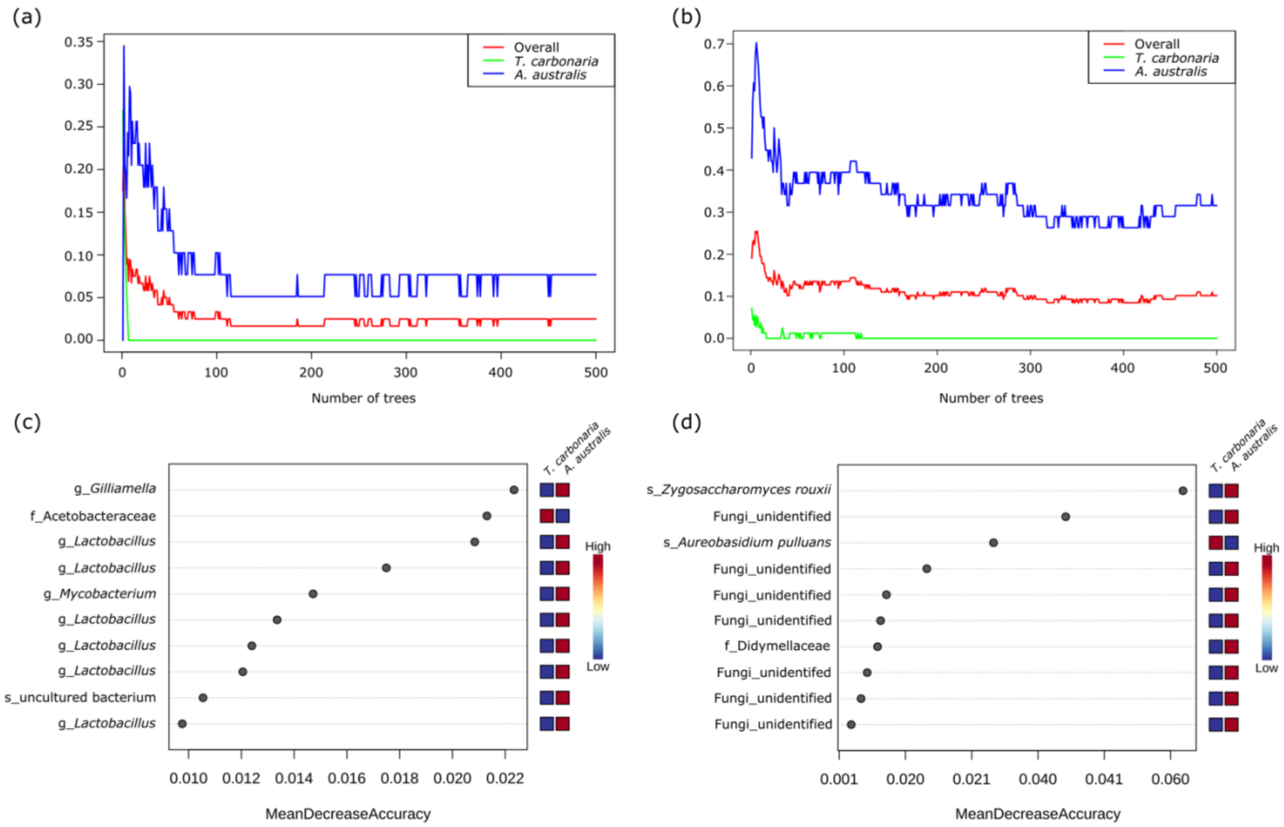


**Fig.S2 Predication of key microbial taxa distinguishing *T. carbonaria* and *A. australis* gut microbiomes using random forest tests**. Changes in error rate with the number of trees shown for the bacterial (a) and fungal community (b). The algorithm achieved stable prediction using >100 trees. Key microbial taxa that are distinct in relative abundance between the two stingless bee species are shown for bacterial (c) and fungal (d) communities.

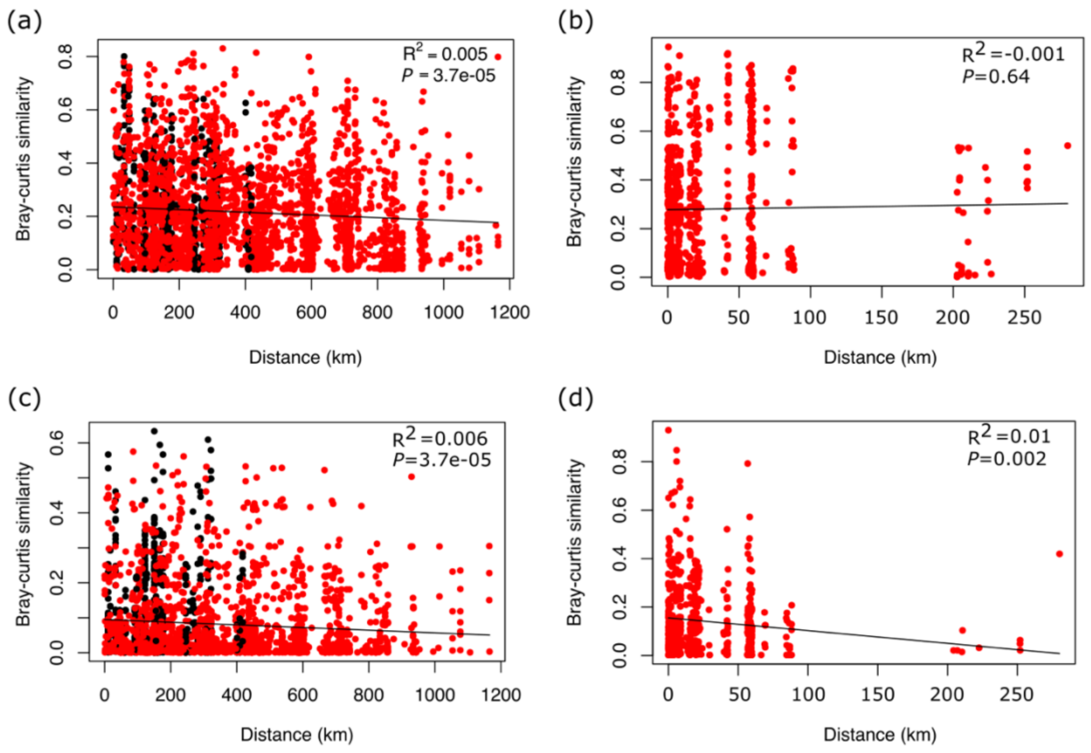


**Fig.S3 Distance-decay in similarity between stingless bee gut microbiomes based on Bray-Curtis similarity**. Bray-Curtis similarity that emphasizes abundant taxa was calculated based on bacterial/fungal OTU tables, and plotted against the geographical distance (Km) for *T. carbonaria* bacterial communities (a), *A. australis* bacterial communities (b), *T. carbonaria* fungal communities (c), and *A. australis* fungal communities (d). Adjusted $R^2$ and *P* values displayed for each linear regression. Wild bee samples (red dots) and managed bee samples (black dots) and regression lines (solid line) shown.

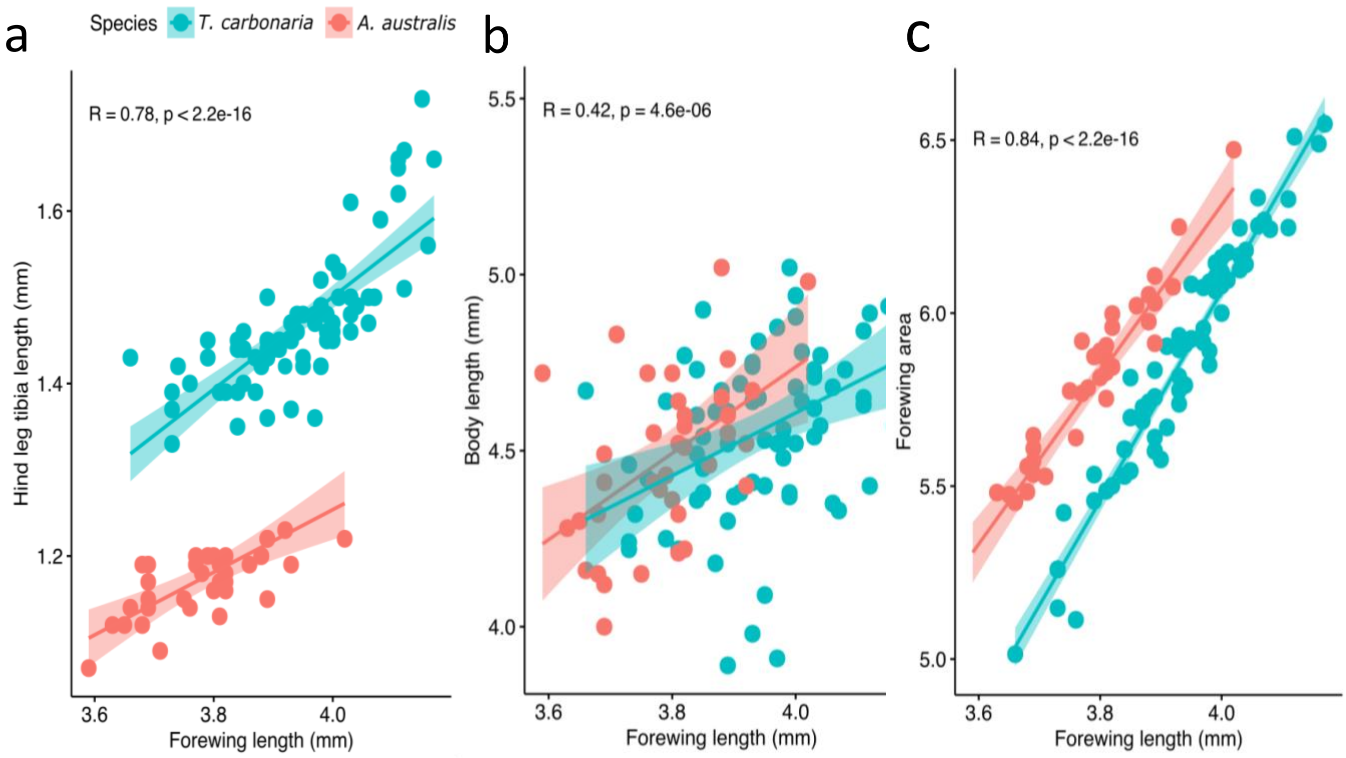


**Fig.S4 Correlation between forewing length and hind leg tibia length (a), full body length (b) and forewing area (c) in *Tetragonula carbonaria* and *Austroplebeia australis* foragers.** Regression lines (solid line), 95% confidence interval (shaded area) and data points (dots) shown.

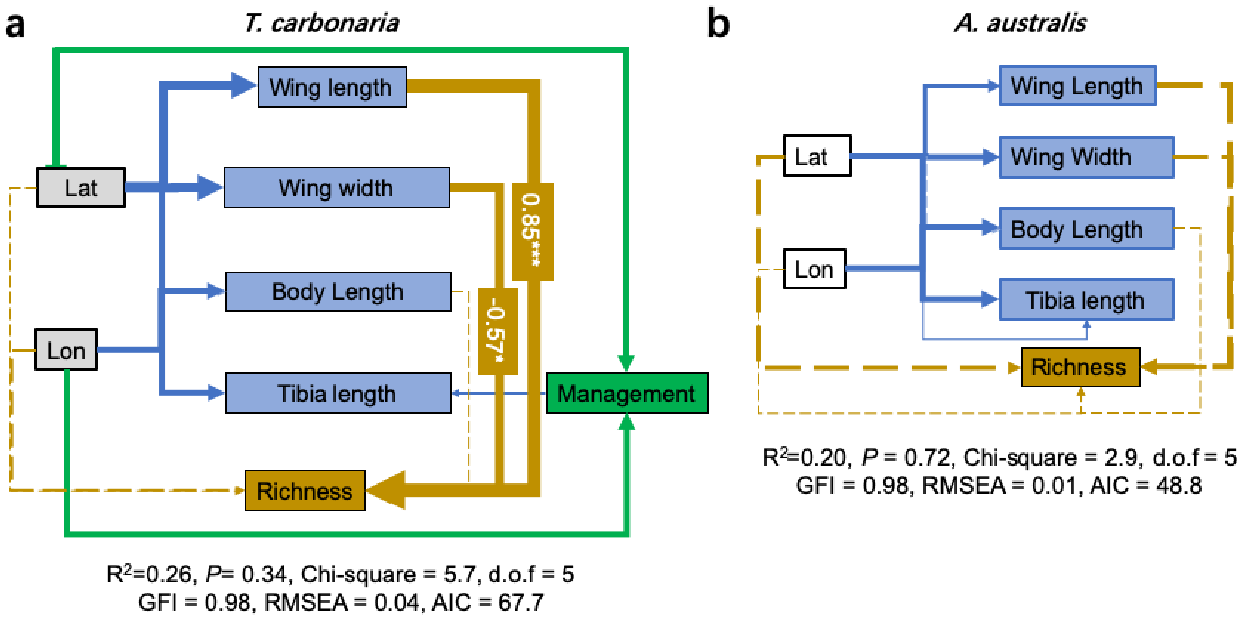


**Fig.S5** Structural equation model (SEM) summarizing correlations of stingless bee morphometric traits, geographical factors and management approaches with the host gut bacterial richness for *Tetragonula carbonaria* (a) and *Austroplebeia australis* (b). Solid arrows indicate significant effect sizes ($P < 0.05$, dashed lines $P > 0.05$) and width of the arrow represents the strength of the relationship. The colour of the arrows correspond to each targeted factor. Standardized path coefficient values are shown besides the significant pathways. Lon: longitude; Lat: latitude.
